# Activation of the *Androgen Receptor* gene by BORIS/CTCFL in prostate cancer cells

**DOI:** 10.1101/195875

**Authors:** Yukti Hari-Gupta, Georgia-Xanthi Kita, Dawn Farrar, Elena Klenova

## Abstract

BORIS/CTCFL, a paralogue of the chromatin architectural protein CTCF, is a member of the cancer-testis antigen family, normally present in the testes. BORIS is expressed in various tumours, including prostate cancers, however the function of BORIS in cancer cells is not well defined. The androgen receptor (AR) plays a critical role in the normal development of a human prostate gland and pathogenesis of prostate cancer. In our previous study we described a positive correlation between elevated levels of BORIS and AR in prostate cancers, and activation of the *AR* gene by BORIS in prostate cancer cells. Elucidation of the mechanisms involved in the modulation of AR activity is important to understand prostate tumourigenesis and investigation of transcriptional regulation of the *AR* gene by BORIS may provide new insights into this issue. Here we report the ability of BORIS to not only positively regulate *AR* in androgen-dependent prostate cancer (ADPC) cells, but re-activate epigenetically silenced *AR* in androgen-independent prostate cancer (AIPC) cells leading to the production of biologically active AR protein. CTCF, on the other hand, had repressive effects on the *AR*. In both, ADPC and AIPC cells, introduction of ectopic BORIS was associated with the reduction in the *AR* promoter methylation, increase in active and decrease in repressive chromatin marks, and decrease in CTCF occupancies at the two main upstream BORIS/CTCF binding sites. We propose a model of epigenetic regulation of *AR* by BORIS in prostate cells whereby BORIS remodels the chromatin at the *AR* promoter leading to transcriptional activation.

## Introduction

BORIS (<u>B</u>rother <u>o</u>f the <u>R</u>egulator of <u>I</u>mprinted <u>S</u>ites), also called CTCFL for “CTCF-like”, is a paralogue of the transcription factor, CTCF[1-4], a ubiquitous protein playing important roles in the regulation of transcription [5, 6], chromatin insulation [7, 8] and global nuclear organization [9-11]. The N- and C-terminal domains of BORIS and CTCF are distinct, whereas the central 11 zinc-finger DNA binding domain in these proteins is almost identical [1, 2]. It is therefore expected that BORIS and CTCF have the ability to bind to similar sequences supported by the genome-wide ChIP-Seq data [12], however recent studies revealed more complex patterns in BORIS binding [13]. BORIS expression is restricted to the testis [1, 2], but it is aberrantly activated in various human cancers, including prostate cancer [14-19]. BORIS functions have not been fully investigated, but it has been reported that BORIS is important for epigenetic reprogramming occurring in the germ cells during development [1, 2, 12, 20, 21]. Furthermore, epigenetic changes introduced by BORIS, have been reported to cause DNA and histone demethylation, chromatin remodelling and activation of transcription of various genes, including cancer genes [22-24] [25-28]. BORIS has been linked to cell proliferation [29] and cancer stem cells and “stemness” gene expression [30, 31]. However, a defined function of BORIS in cancer cells still remains debatable [17, 32, 33].

Our previous study revealed an association between BORIS and Androgen Receptor (AR) whereby the elevated levels of BORIS in prostate cancer cell lines and tumours not only correlated with high AR levels in prostate cancer cells, but BORIS could stimulate expression of the endogenous *AR* gene [18]. AR is a member of the nuclear receptor superfamily [34] activated by androgens, in particular testosterone, primarily produced by the testes with a small contribution from the adrenal glands [35], which in prostate cells is converted into a more active hormone, dihydrotestosterone (DHT) [36]. In the absence of the ligand, the AR is located primarily in the cytoplasm, however following the interaction of DHT with AR, the ligand-induced receptor translocates to the nucleus where it forms a homodimer and binds to the androgen response elements (AREs) of the androgen-sensitive genes [37]. Many AR target genes have been identified and are known to be involved in the maintenance of prostate growth and development [38-41]. The AR mediates various functions of androgens essential for cell viability, proliferation and invasion [42, 43]. It is also one of the key factors in the pathogenesis of androgen dependent prostate cancer (ADPC), which initially relies on the AR signalling pathways [44, 45] and hence the androgen-ablation therapy has been the most common treatment for ADPC patients [46, 47]. However, ADPCs eventually become androgen-independent and ultimately progress to more aggressive androgen independent prostate cancer (AIPC). The molecular mechanisms of these phenomena are not well understood, however a wide range of genetic and epigenetic mechanisms have been implicated in the development of refraction to androgens [48-50]. Epigenetic changes (e.g. *AR* methylation, chromatin modifications, transcriptional control) are of particular interest as they can be reversed [51, 52]. It is therefore important to investigate the molecular events involved in the epigenetic regulation of the *AR*, in particular the role of transcription factors in these processes. For example, LEF1 and YB-1 were reported to activate [53, 54] and p53 to repress *AR* [55]. Various functional elements have been identified and binding sites for more than 20 transcription factors mapped in the *AR* promoter region [49].

The correlation found between BORIS and AR expression in prostate tumour tissues[18] led to the hypothesis that BORIS can be involved in prostate tumour progression as a direct transcriptional regulator of *AR*. In the present study we demonstrate that BORIS plays an important role in the activation of the *AR* gene in ADPC and AIPC cells and proposed a model of epigenetic regulation of *AR* by BORIS.

## Materials and Methods

### Cell Lines

Prostate cell lines used in this study were the AR-positive and androgen dependent prostate cancer (ADPC) cells, LNCaP and VCaP; AR-negative and androgen independent prostate cancer (AIPC) cells DU145, partially AR-positive and androgen independent prostate cancer (AIPC) cells, PC3, and AR-negative and androgen independent benign prostate cells, BPH-1. Cells were maintained in RPMI 1640 medium supplemented with L-Glutamine (PAA-GE Healthcare), 50μg/ml Gentamicin (Life Technologies-Invitrogen) and 10% foetal bovine serum (Biosera).

### Transient transfections with plasmid DNA and siRNA

The expression vectors (pCMV6-BORIS, pCI-CTCF, pEGFP-BORIS) used in transient transfections have been previously described. Transient transfections were performed using a calcium phosphate transfection protocol [56]. Hs_CTCF_4 siRNA (Qiagen, Manchester, UK), BORIS SMARTpool siRNA (Dharmacon, Epsom, UK) and non-target siRNA (Dharmacon, Epsom, UK) were used for transfections at a concentration of 50 pM. Cells were seeded at a density of 1.2 × 10^5^ and transfected on the following day with siRNA and DharmaFECT2 (Dharmacon) according to the manufacturer’s protocol.

### Western-blot analysis

For Western blot analysis, lysates from cells were prepared according to Klenova et al [57] with modifications. The primary antibodies were used at the following dilutions: mouse anti-BORIS monoclonal antibody (clone A47, MABE 1125, Millipore), 1:1000; mouse anti-Androgen Receptor (AR) antibody (AR441, Abcam), 1:100; the mouse anti-CTCF antibody (BD Biosciences, Oxford, United Kingdom), 1:500; the rabbit anti-His tag (Cell Signalling), 1:1000; the mouse anti-α-tubulin (Sigma) antibody, 1:2000. The secondary antibodies were as follows: Goat anti-mouse IgM-HRP (Abcam) (used at 1:10,000 dilution), Goat anti-mouse IgG-HRP (Abcam) (used at 1:10,000 dilution) and Goat anti-rabbit IgG-HRP (Abcam) (used at 1:15,000 dilution). Detection was performed with enhanced chemiluminescence reagent (Interchim) according to the manufacturer’s instructions.

### Immunofluorescence staining

Immunofluorescence staining was performed on prostate cancer cells transiently transfected with pEGFP-BORIS expression vector as previously described followed by incubation with 100 nmol of DHT (5α-Androstan-17β-ol-3-one) (SIGMA). After 24 hours of DHT treatment, cells were fixed with 4% paraformaldehyde, permeabilized with 0.25% Triton X-100 in 1xPBS, blocked with 2% normal goat serum in 1xPBS: 0.05% Tween: 1% BSA, incubated with the mouse anti-AR antibody (AR441-Abcam; Ab dilution 1:100) and then with the TRITC- labelled anti-mouse IgG antibody (Southern Biotech, Antibody dilution 1:200). The cells were mounted on the microscope slides with Fluoro-Gel mounting media containing DAPI (4’,6-diamidino-2-phenylindole dilactate) (Interchim). Images were taken at x60 magnification using Nikon Eclipse Ti microscope. Staining of cells expressing EGFP-BORIS without DHT exposure was also performed to visualize the differences in AR localization under the effect of DHT.

### Chromatin Immunoprecipitation Assays

Chromatin immunoprecipitation (ChIP) assays were performed from cross-linked cells (1 × 10^6^) using Protein A agarose/salmon sperm DNA beads (Millipore) for rabbit (IgG) polyclonal anti-CTCF antibody (Millipore), rabbit polyclonal anti-H3K4me3 (active chromatin) (Millipore), rabbit polyclonal anti-H3K27me3 (repressive chromatin) (Millipore) and Protein L agarose beads (Thermo Scientific) for mouse monoclonal (IgM) anti-BORIS N-terminal antibody. Protein L Agarose beads were treated with 1 μg sheared salmon sperm DNA (Ambion) according to the manufacturer’s instructions. Cross-linking and immunoprecipitation procedures were followed as described before. Immunoprecipitated DNA was extracted with phenol/chloroform and ethanol precipitated. Real-time PCR was carried out in triplicate using 2 μl of the immunoprecipitated DNA sample and input DNA and 300 nM primers diluted to a final volume of 25 μl in SensiMix Plus SYBR (Quantace). Percentage of DNA brought down by ChIP (% input) was calculated as follows: input = AE^(Ct_input_ – Ct_ChIP_) × Fd × 100% (AE is amplification efficiency, Ct_ChIP_ and Ct_Input_ are threshold values obtained from exponential phase of qPCR, and Fd is a dilution factor of the input DNA to balance the difference in amounts of ChIP). Primers and conditions for qPCR are described in Supplemental Table 1.

### Purification of RNA, Reverse Transcription, and Real-Time Reverse Transcription-Quantitative Polymerase Chain Reaction

Total RNA was isolated with Trisure (Bioline, London, United Kingdom) as described by the manufacturer. RNA was treated with TURBO DNA-free kit (Ambion, Paisley, United Kingdom); 1 μg of total RNA was used for reverse transcription (RT) with VERSO cDNA kit (Thermo Fisher, Loughborough, United Kingdom). For RT-quantitative polymerase chain reaction (qPCR), cDNA samples were diluted at 1:10. A 25-μl PCR mix consisted of 1x SensiMix Plus SYBR (Quantace, London, United Kingdom), 2 μl of diluted cDNA, 200 nM of each primer, and 3 mM MgCl_2_. Amplification, data acquisition, and analysis were carried out using the Bio-Rad CFX Real-Time PCR (Bio-Rad Laboratories). The comparative *C*_t_ method was used to assess relative changes in *mRNA* levels [58]; calculations were made according to Pfaffl [58]. Primers and conditions for RT-qPCR are described in the Supplemental Table 2.

### DNA methylation analysis using methylation dependent restriction enzyme McrBC

A restriction enzyme, McrBC (NEB) was used to analyze DNA methylation within the *AR* promoter. It specifically cleaves DNA containing methyl cytosine preceded by a purine nucleotide (A or G) [59, 60]. The reaction consisted of 500ng of genomic DNA, 5 units of McrBC enzyme, 1x NEB buffer 2, 1x BSA, 1mM GTP and water in a total reaction volume of 50μl. The reaction was incubated at 37°C for 16 hrs followed by PCR performed on digested and undigested DNA, using the *AR* promoter specific primers as follows FP: 5’ CTTGGTCATGGCTTGCTCCT 3’ and RP: 5’AGGACGATAGGAATACCTGCTC 3’. Quantification of the PCR bands was performed using ImageJ software and the amount of methylated DNA was calculated by subtracting DNA left after digestion with McrBC from the total DNA before digestion: the percentage was calculated according to the formula: % Methylation = [ (-McrBC) – (+McrBC)] × 100/ (-McrBC).

### Statistical Analysis

Statistical analysis was carried out using unpaired Student’s T-test. A significant value was detected when the probability was below the 5% confidence level (*P* < .05).

## Results

### AR gene and protein expression analysis in prostate cancer cells with manipulated levels of BORIS and CTCF

We previously reported that expression from the endogenous *AR* gene increased in the AR positive and androgen dependent prostate cancer (ADPC) cells, LNCaP, transiently transfected with the plasmid containing *BORIS* cDNA [18]. We extended this analysis by examining whether BORIS will be able to activate *AR* in the AR-negative and androgen independent prostate cancer (AIPC) cells, DU145. We also aimed to assess the effects of a BORIS paralogue, CTCF, on *AR* since both, CTCF and BORIS, are present in prostate cancer cells. For these experiments, the levels of BORIS and CTCF in LNCaP and DU145 cells were transiently increased or decreased, using expressing vectors or siRNAs, respectively. Efficient knockdown of CTCF and BORIS was achieved following optimisation of transfection with the corresponding siRNAs and siGLO cyclophilin B siRNA (Supplemental Figure 1). In agreement with our previous findings [18], *AR* mRNA expression in LNCaP cells increased in cells over-expressing BORIS and decreased in cells depleted of BORIS. Over-expression of BORIS was associated with the increase in the *AR* mRNA levels in DU145 cells, although to a much lesser extent than in LNCaP cells and no change was detected in cells with reduced BORIS levels (Figure 1A, left panel). BORIS over-expression and down-regulation in both cell lines was confirmed using RT-qPCR (Figure 1A, right panel).

**Figure 1.**
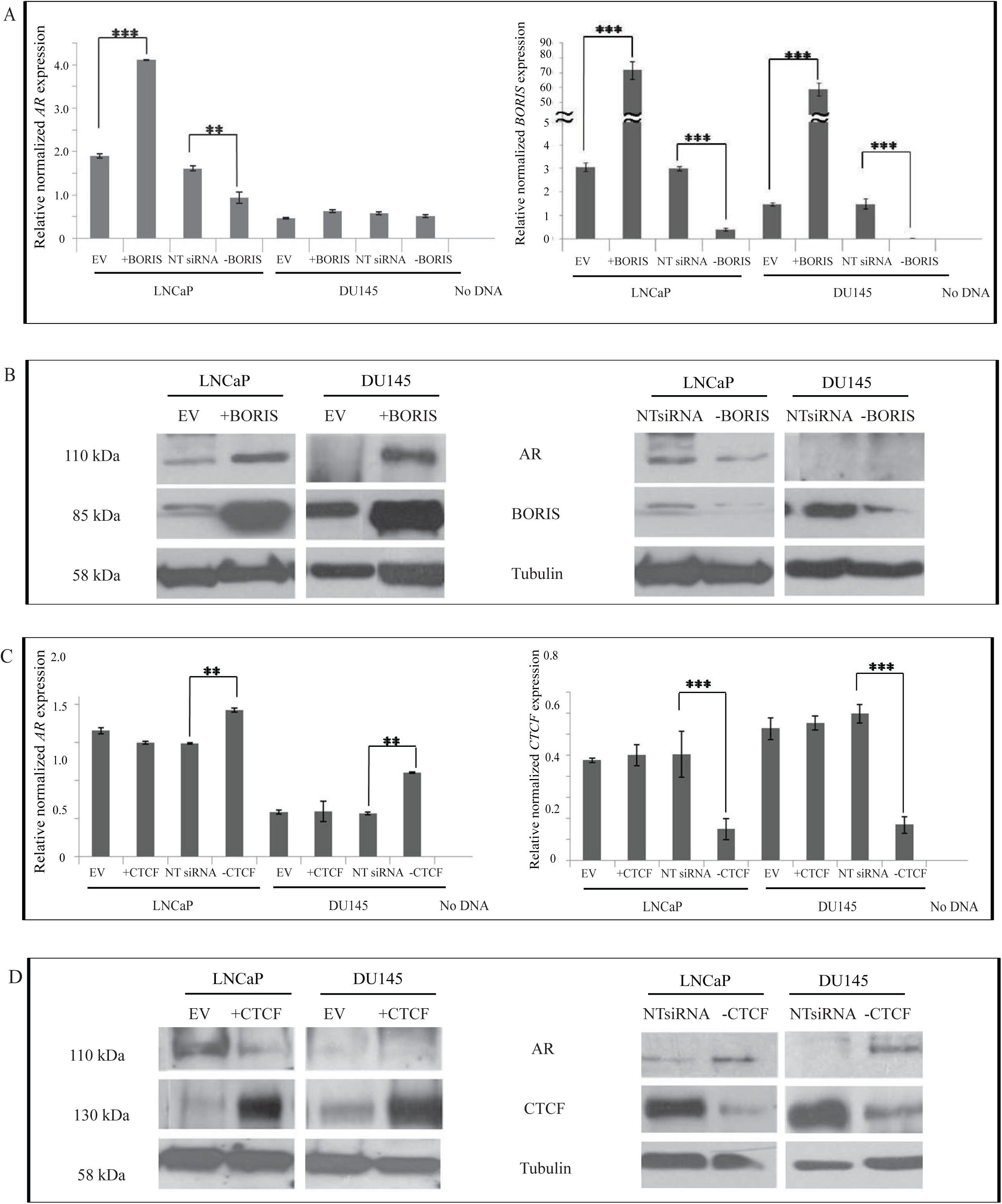
Assessment of the androgen receptor (*AR*) gene and AR protein expression in LNCaP and DU145 cells with transiently increased or decreased levels of BORIS and CTCF. Cells were transfected with 1 μg of empty vector (EV), pCMV6-BORIS (+BORIS), pCI-CTCF (+CTCF) or 100pm of non-target (NT) siRNA, BORIS siRNA (-BORIS) or CTCF siRNA(-CTCF) and harvested 48 hours post-transfection. Total RNA was prepared and analyzed by RT-qPCR. The expression levels of *BORIS, CTCF* and *AR* mRNA were calculated using the comparative *C*_t_ method (δδ*C*_t_) and normalized to *β-actin* mRNA expression used as the reference gene. Each data point is an average of three independent experiments and error bars indicate the standard deviation (Student’s T test: ** P<0.01; *** P<0.005). For Western blot analysis, cells lysates were obtained, 20μg total protein were loaded onto 8.1% SDS-PAGE blotted and probed with relevant antibodies to assess the levels of CTCF (130 kDa), BORIS (85 kDa) and AR (110 kDa); α-tubulin antibody was used as loading control. **A.** The *AR* (left panel) and *BORIS* (right panel) mRNA levels in LNCaP and DU145 cells with modulated levels of BORIS. **B.** AR and BORIS protein levels in LNCaP and DU145 cells transiently transfected with empty vector (EV) and pCMV6-BORIS (+BORIS) (left panel) and non-target (NT) siRNA and BORIS siRNA (-BORIS) (right panel). **C.** The *AR* (left panel) and *CTCF* (right panel) mRNA levels in LNCaP and DU145 cells with modulated levels of CTCF. **D.** AR and CTCF protein levels in LNCaP and DU145 cells transiently transfected with empty vector (EV) and pCI-CTCF (+CTCF) (left panel) and non-target (NT) siRNA and CTCF siRNA (-CTCF) (right panel).

Reduction in CTCF caused up-regulation in the *AR* mRNA levels, i.e. the effects opposite to BORIS depletion in both cell lines. However, very little or no difference in *AR* expression was observed in samples with high levels of CTCF (Figure 1C, left panel). The changes in *CTCF* mRNA in cells depleted of CTCF were significant, however, only a very small change in the *CTCF* mRNA levels were detected in samples transfected with pCI-CTCF (Figure 1C, right panel). This is surprising given that the ectopic CTCF protein was present at high levels (see below).

Next, the assessment of AR, CTCF and BORIS expression at the protein levels was carried out. As shown in Figure 1B (left panel), both the LNCaP and DU145 cells over-expressing BORIS produced increased amounts of the AR protein. In LNCaP cells depleted for BORIS, the AR protein levels were decreased. Together with the RT-qPCR data, these results suggest AR re-activation in BORIS-over expressing DU145 cells. It should be noted that the accumulation of BORIS is higher at the protein level than at the RNA level. However, their comparisons may not be accurate in some cases where, for example, mRNA half life could be much shorter whereas protein stability higher (this is further considered in the Discussion section).

The amounts of the AR protein were also found to be affected by CTCF in both LNCaP and DU145 cells, whereby depletion from CTCF resulted in the increase of the AR protein and vice versa (Figure 1D). Notably, although the AR protein was not present in the control DU145 cells, it appeared in cells depleted from CTCF. These effects were more pronounced at the protein level, which implies that in these cells *CTCF* mRNA may be much less stable than the CTCF protein.

In summary, the analysis of the *AR* expression at the mRNA as well as protein levels suggest that *AR* is positively regulated by BORIS and negatively regulated by CTCF, although the latter was demonstrated only in CTCF knock down experiments. The possibility of re-expression of AR in DU145 (AIPC) cells is significant because it may be exploited in the treatment of hormone resistant prostate cancers.

### Immunofluorescence analysis of AR in ADPC and AIPC cells transfected with pBORIS-EGFP plasmid DNA in the presence and absence of DHT

The aim of our next experiments was to investigate whether the introduction of the ectopic BORIS would result in the production of nuclear AR upon androgen stimulation, which is one of the prerequisites for AR protein to function as the ligand-dependent transcription factor [38]. The response to DHT in LNCaP and DU145 cells was tested prior to these experiments demonstrating, as expected, the increased levels of AR in LNCaP and no AR induction in DU145 cells (Supplemental Figure 2). Additionally, the functionality, i.e. the ability of the EGFP-BORIS fusion protein to up-regulate AR, was confirmed in several cell lines, ADCP and AICP (Supplemental Figure 3 and data not shown). Following transfection with plasmids expressing EGFP-BORIS fusion and the EGFP protein only, the ADPC prostate cancer cells, LNCaP, and AIPC cells, PC3 and DU145, were treated with DHT, and BORIS and AR were visualised by immunofluorescent microscopy. In LNCaP cells, AR was found mostly in the cytoplasm in the absence of DHT. Clear nuclear staining of AR was observed in the presence of DHT in cells both non-transfected and transfected with pEGFP-BORIS, although the signal from the latter was stronger (Figure 2A, panel i). In control LNCaP cells transfected with EGFP empty vector, AR was mostly present in the cytoplasm in the absence of DHT and in the nucleus after addition of DHT (Figure 2B, panel i).

**Figure 2.**
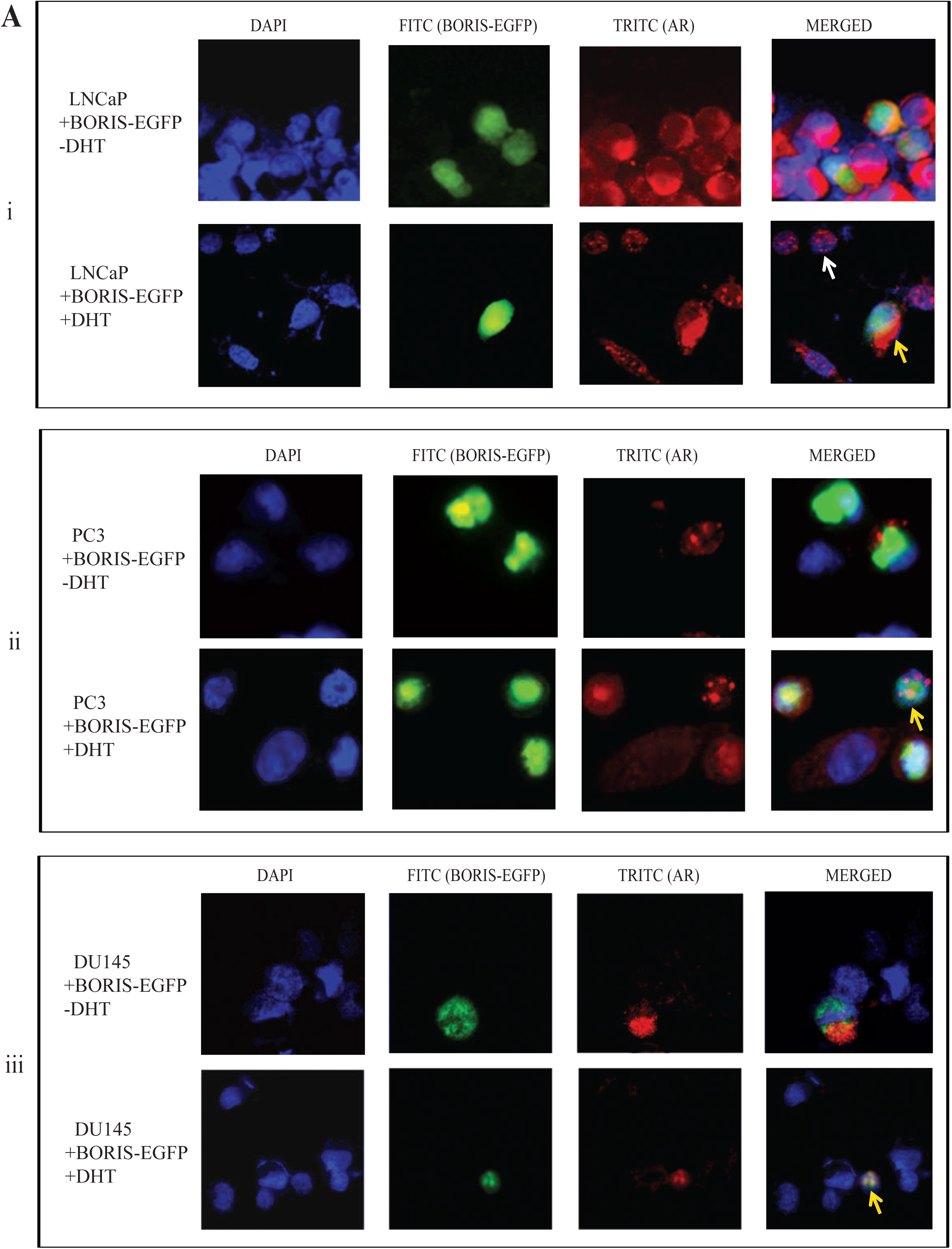
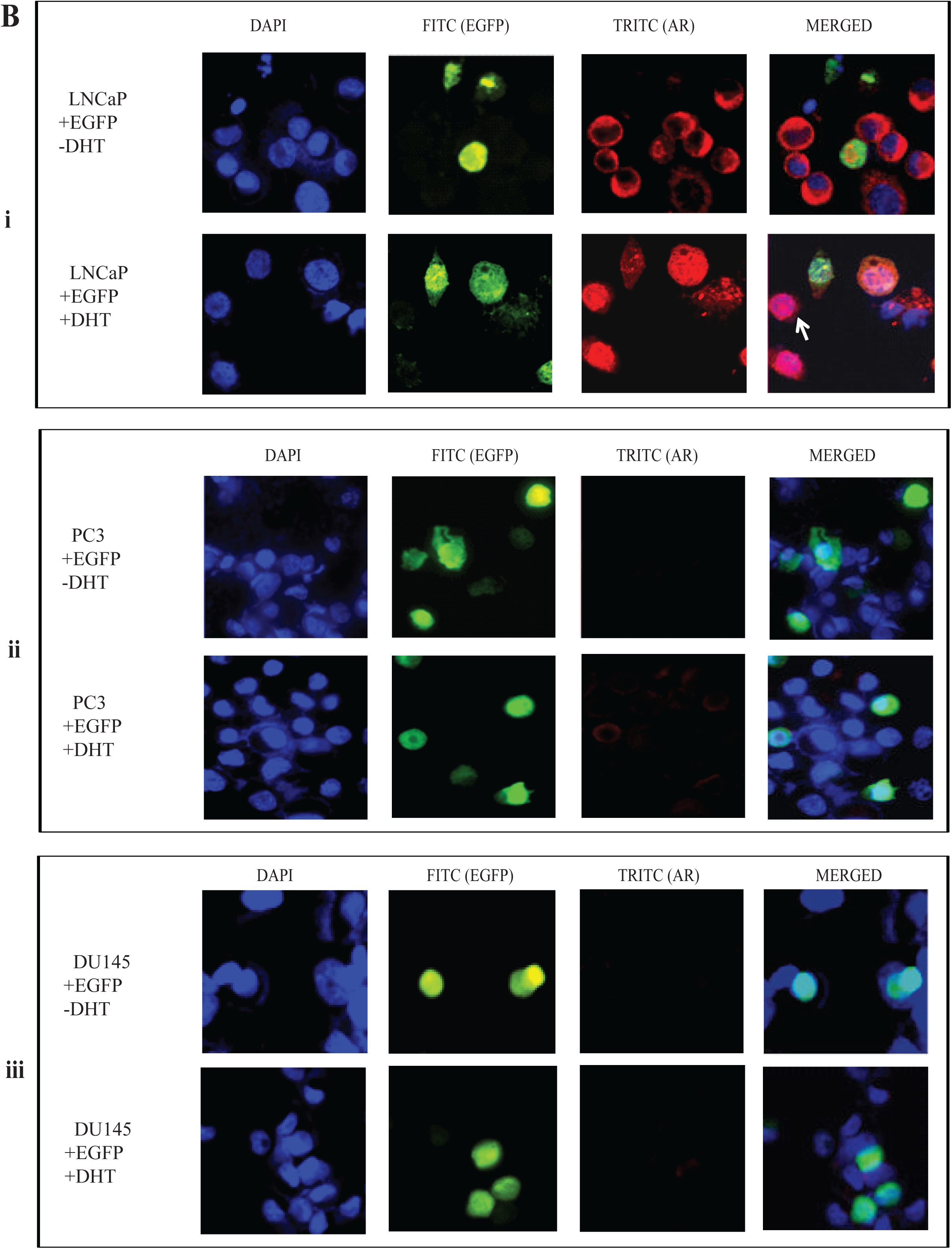
Immunofluorescence analysis of AR expression in LNCaP, PC3 and DU145 cells transiently transfected with pBORIS-EGFP in the presence and absence of androgen (DHT). LNCaP (section i), PC3 (section ii) and DU145 (section iii) cells (1.5x10^5^) were transfected with 1μg of pBORIS-EGFP (panel **A**) or pEGFP (panel **B**) plasmids and AR distribution in cells was investigated in the presence (+) or absence (-) of DHT. Transfected cells were identified using the FITC (green) filter and the AR expression was detected by the staining with the primary anti-AR monoclonal antibody followed by TRITC (red) labeled secondary mouse antibody. Nuclei were visualized with 4’,6- diamidino-2-phenylindole (DAPI) staining (*blue*). *Right*, merge of the green and red channels. Images were taken at x60 magnification using Nikon Eclipse Ti microscope. Examples of LNCaP cells demonstrating nuclear localization of AR in the presence of DHT are indicated by white arrows. Examples of LNCaP, PC3 and DU145 cells containing ectopic BORIS and demonstrating nuclear localization of AR in the presence of DHT are indicated by yellow arrows.

**Figure 3.**
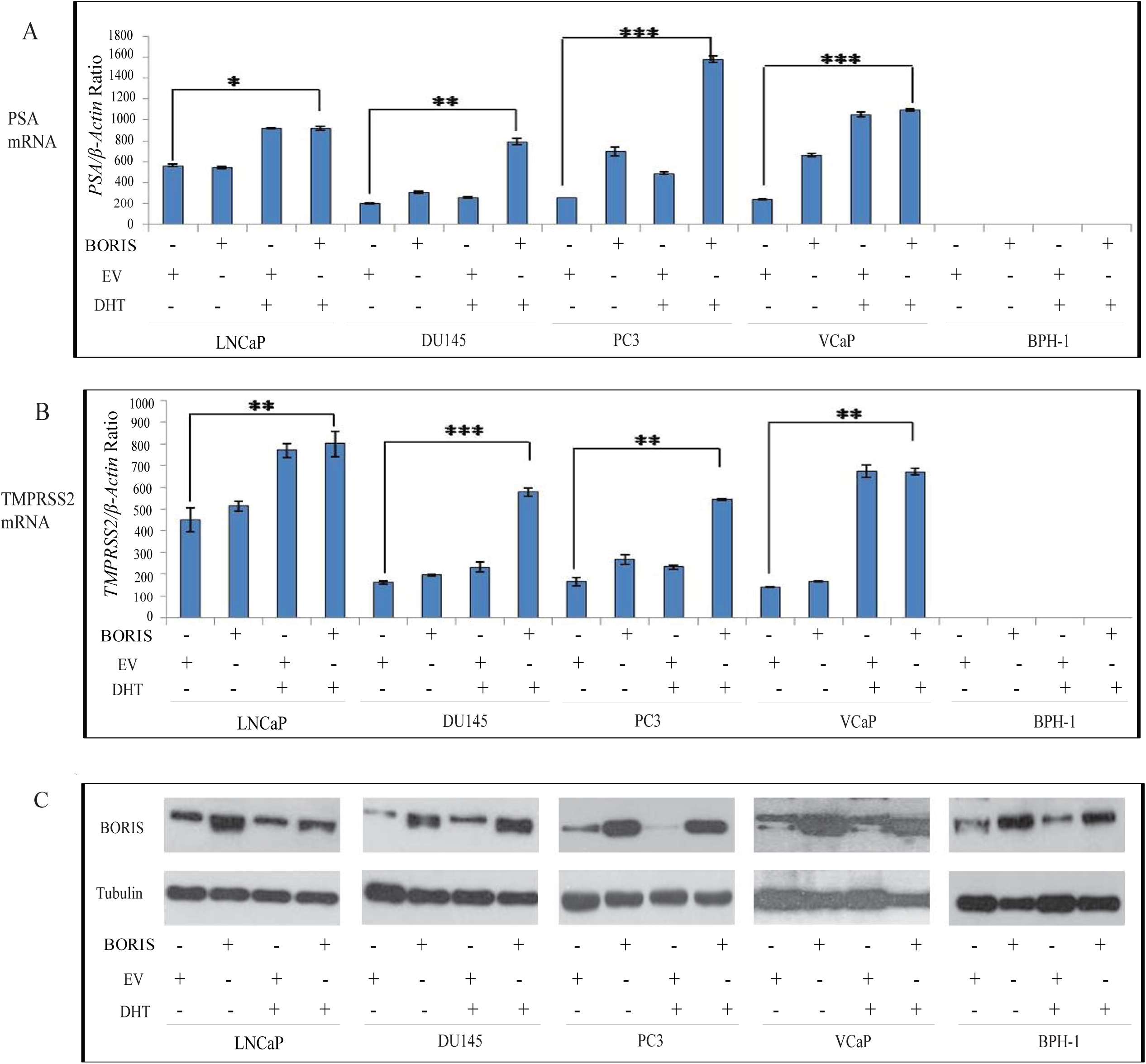
Analysis of *PSA* and *TMPRSS2* gene expression as a measure of AR activity. Five different prostate cancer cell lines were transfected with 1μg of pCMV6-BORIS DNA or empty vector (pEV) followed by 24 h of DHT treatment as indicated by (+) or (-). Control cells with no DHT treatment (-) were incubated with 100% ethanol (equal volume as DHT) for 24 hrs. Cells were then harvested and RNA and protein lysates prepared. RT-qPCR using specific primers was carried out for quantification of *PSA* **(A)** and *TMPRSS2* **(B)** mRNA. The results for each mRNA were normalized using *β-Actin* mRNA used as the reference gene. Each data point is an average of three independent experiments and error bars indicate the standard deviation (Student’s T test: * P<0.05; ** P<0.01; *** P<0.005). **C.** Western blotting was performed to confirm BORIS over-expression in prostate cells transfected with pCMV6-BORIS or empty vector (pEV) in the presence (+) and absence (-) of DHT; α- tubulin was used as loading control.

Introduction of the ectopic BORIS into PC3 and DU145 cells resulted in the appearance of AR in these cells and following treatment with DHT, the characteristic for AR nuclear staining was observed (Figure 2A, panels ii and iii). The merged images from bottom panels of PC3 and DU145 cells demonstrated some co-localization of AR and BORIS-EGFP in the nucleus (yellow dots). No AR was detected in the control (pEGFP transfected) PC3 and DU145 AIPC cells, treated and untreated with DHT (Figure 2B, panels ii and iii). These results further supported the role of BORIS in stimulating AR expression in AIPC; the re-location of AR to the nucleus in the presence of DHT implied that the induced AR is functional.

### Analysis of expression from the AR-responsive genes in prostate cancer cells containing ectopic BORIS

To further confirm AR functionality, expression levels of genes regulated by (*PSA* and *TMPRSS2*) in BORIS-over expressing ADPC and AIPC cells were analyzed. Prostate Specific Antigen (*PSA*) and Transmembrane protease, serine 2 (*TMPRSS2*) are well characterized AR target genes associated with prostate cancer growth[61] and are known to be up-regulated by AR in the presence of DHT in ADPC cells [62-64]. On the other hand, very low or no expression of *PSA* and *TMPRSS2* have been reported in AIPC cells such as PC3 and DU145 that can be explained by the lack of functional AR in these cells [63, 65]. In our experimental context, these genes represent suitable candidates to assess the androgen-dependent AR activity.

A panel of cell lines used in these experiments included two ADPC (VCaP, LNCaP), two AIPC (PC3, DU145) and AR-negative benign prostate hyperplasia (BPH-1) cells. As expected, the ADPC were positive for *AR* mRNA and the appearance of the *PSA* and *TMPRSS2* mRNA was observed in these cells treated with DHT, whereas three other cell types cells were *AR*, the *PSA* and *TMPRSS2* mRNA negative (Supplemental Figures 4 and 5). In order to test the effects of BORIS on *PSA* and *TMPRSS2* expression, ADPC and AIPC cells were transfected with pCMV6-BORIS and the empty vector (EV) and then treated with DHT. As shown in Figure 3C, BORIS protein was over-expressed in all cells transfected with pCMV6-BORIS, In agreement with published reports, *PSA* and *TMPRSS2* expression in ADPC cells VCaP and LNCaP increased after treatment with DHT [63, 64], with and without ectopic BORIS. Interestingly, in AIPC cells, DU145 and PC3, but not BPH-1, in the presence of ectopic BORIS and DHT, the levels of *PSA* and *TMPRSS2* mRNA reached levels comparable with those in the ADPC cells (Figures 3A and 3B).

**Figure 4.**
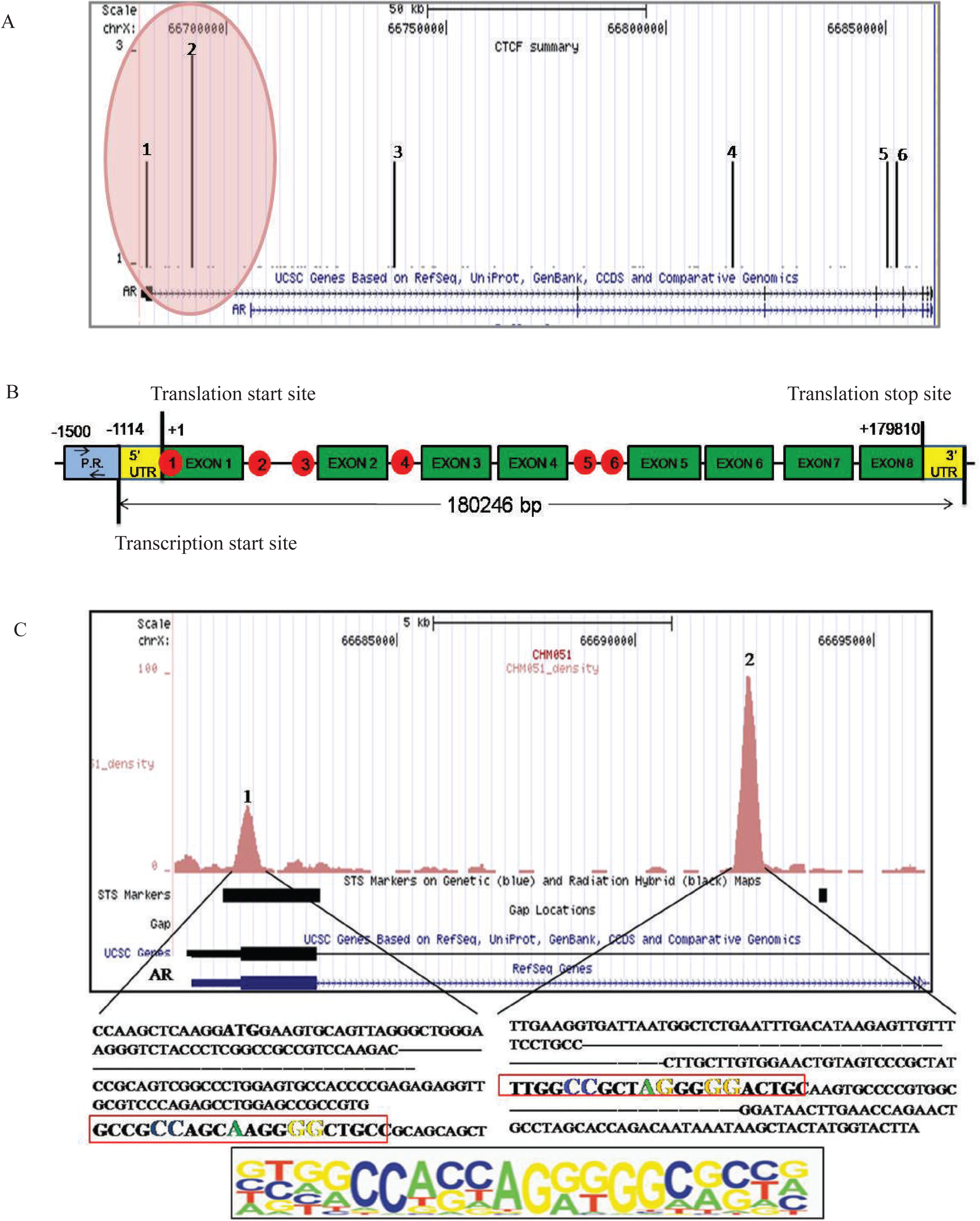
Representation of CTCF target sites (CTSs) within the human *AR* gene. **A.** Six potential CTCF target sites (CTS) were mapped within the human *AR* gene using UCSC genome browser http://genome.ucsc.edu/ Feb 2009 (GRCh37/hg19). **B.** A schematic representation of the location of all *AR* CTSs within the human *AR* gene. Exons are shown as green boxes and *AR* CTSs are presented as red circles. *AR* CTS-1 is positioned near the translational start site and continues into exon 1.The *AR* CTS-2 through to CTS-6 are present in the intronic regions. Promoter region (P.R.) is shown in blue (-1114 to -1500), downstream of the 5’UTR. Arrows depict promoter specific primers used in promoter methylation studies. The coordinates of CTSs 1 to 6 in the *AR* gene (from the translation start site) are: Site1: -12 to 388; Site 2: 10391-10780; Site 3: 56389-56788; Site 4: 133589-133988; Site 5: 168789-169189; Site 6: 170789-171188. **C.** The ChIP data are zoomed in and the sequences for *AR* CTS-1 and CTS-2 are expanded. Both sites contain CTCF consensus motif (shown in large coloured letters at the bottom of panel C). The conserved motif within the *AR* CTS-1 and CTS-2 sequence is outlined in red.

**Figure 5.**
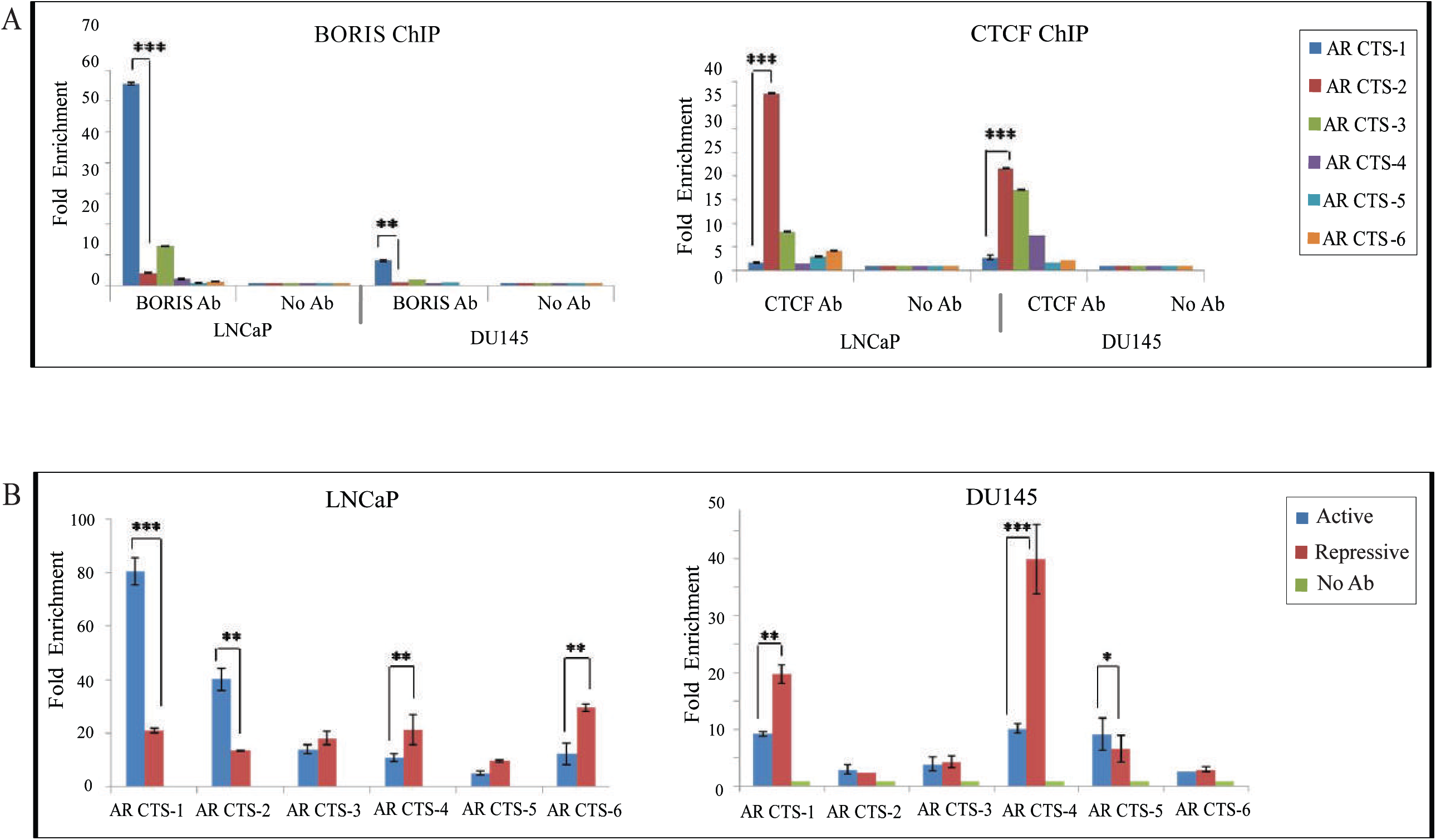
Analysis of BORIS and CTCF binding and distribution of chromatin marks at the *AR* CTSs at the six predicted *AR* CTSs in LNCaP and DU145 cells. **A.** BORIS ChIP (left) and CTCF ChIP (right). DNA obtained after immunoprecipitation with the anti-BORIS or anti-CTCF antibodies was analyzed by qPCR using primers for the *AR* CTS-1, 2, 3, 4, 5, 6 and the results were calculated as the percentage of input chromatin precipitated at the region of interest. The DNA was analyzed for *AR* CTS-1 through to CTS-6 by qPCR using corresponding primers; all reactions were performed in triplicates and mean fold enrichment relative to the control ChIP experiment with no antibody (designated as 1.0) were calculated; error bars indicate the standard deviation (Student’s T test comparing binding between *AR* CTS-1 and *AR* CTS-2: ** P<0.01; *** P<0.005). **B.** ChIP was performed using the antibodies to identify active (anti-H3K4me3) and repressive (anti-H3K27me3) chromatin marks in two prostate cancer cell lines, LNCaP (left) and DU145 (right). The ChIP DNA samples were analyzed by qPCR using primers for amplification of the *AR* CTS-1 through to CTS-6. The DNA was analyzed for *AR* CTS-1 through to CTS-6 by qPCR using corresponding primers; all reactions were performed in triplicates and mean fold enrichment relative to the control ChIP experiment with no antibody (designated as 1.0) were calculated; error bars indicate the standard deviation (Student’s T test comparing binding between *AR* CTS-1 and *AR* CTS-2: * P<0.05; ** P<0.01; *** P<0.005).

Our results suggest that BORIS over expression not only restored AR expression in AIPC cells but also produced a functional AR that was able to regulate expression of target genes in the presence of DHT. However no *PSA* and *TMPRSS2* mRNAs were observed in BORIS-over expressing BPH-1 cells; this is addressed further in the Discussion section of this paper.

### Analysis of the ChIP-seq data for CTCF binding in the human AR gene using the UCSC genome browser

The *AR* gene in DU145 and PC3 is known to be silenced due to promoter hypermethylation [66, 67]. We hypothesized that the *AR* re-activation could be due to epigenetic changes introduced by BORIS, reported to cause DNA and histone demethylation, chromatin remodelling and activation of transcription [22-24] [25-27]. BORIS and CTCF contain a similar DNA binding domain and therefore are able to interact with similar CTCF target sequences (CTSs) and function in a mutually exclusive manner in DNA binding and transcriptional regulation [1, 2]. In order to identify potential CTCF/BORIS binding sites in the *AR* gene we interrogated the ChIP-Seq data deposited in the University of California Santa Cruz (UCSC) genome browser [68] (http://genome.ucsc.edu/ Feb 2009 (GRCh37/hg19). Within the human *AR* gene, the enrichment in CTCF binding was detected in six regions, named as *AR* CTS-1 through to CTS-6 (Figure 4). The *AR* CTS-1 is positioned near the translational start site and continues into exon 1, whereas *AR* CTS-2 through to CTS-6 are located within the intronic regions (Figure 4B). The enrichment in CTCF binding was more pronounced at the *AR* CTS-1 and *AR* CTS-2 than at other *AR* CTSs and was observed in a wide range of cell lines deposited in the USCS genome browser (Supplemental Figure 6). The *AR* CTS-1 and *AR* CTS-2 contain sequences that comply with the conserved 20-mer CTCF consensus binding motif described in previous reports [7, 69] (Figure 4C). Due to their ability to bind the same CTSs, the *AR* is likely to be regulated by both, CTCF and BORIS, competing with each other for some or all of the six *AR* CTSs, although the *AR* CTS-1 and *AR* CTS-2 may be more important for the *AR* regulation than other CTS due to their proximity to the *AR* promoter region.

**Figure 6.**
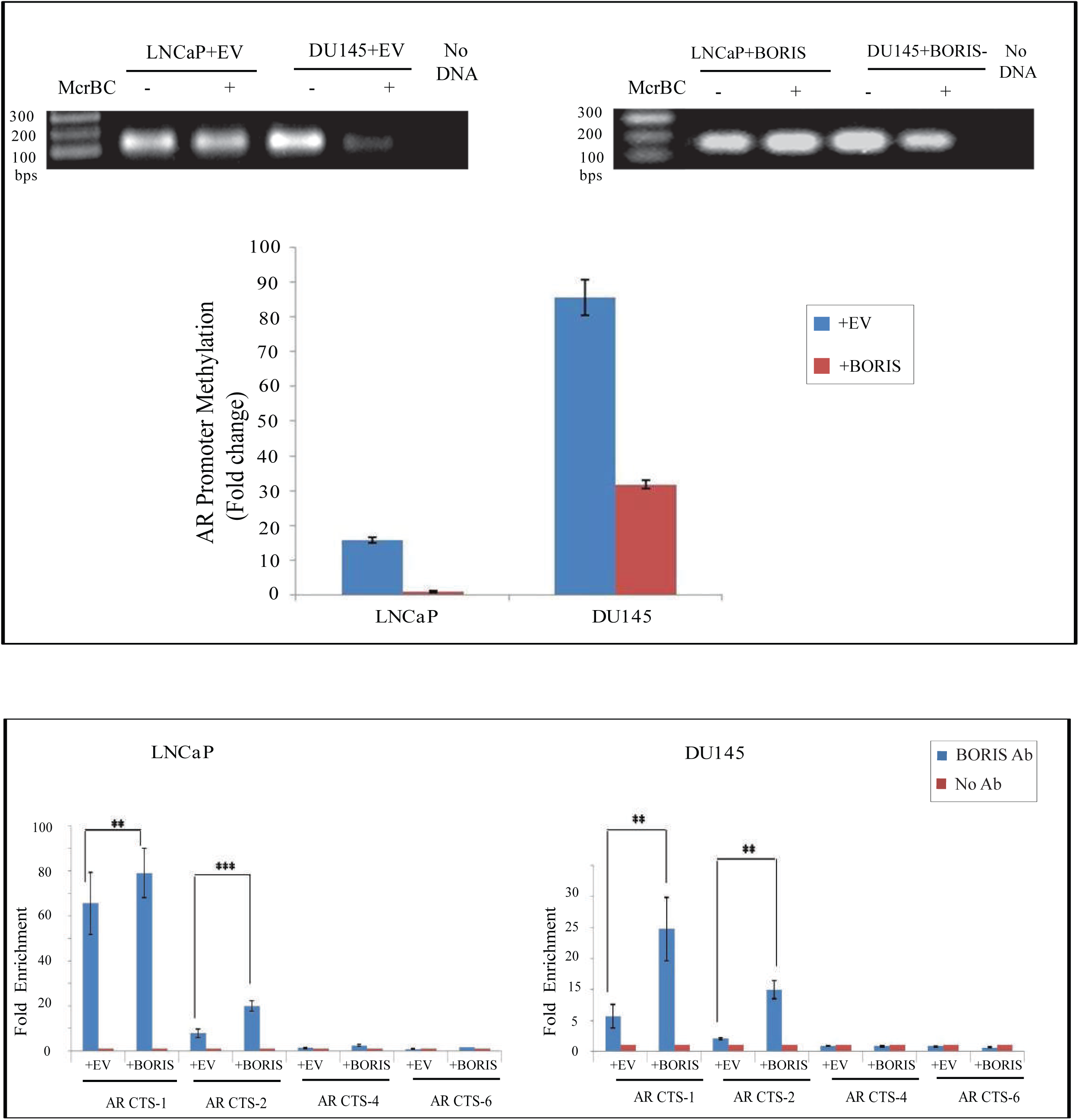
Analysis of methylation of the *AR* promoter in prostate cancer cells over-expressing BORIS, using McrBC enzyme digestion. **A.** Prostate cancer cells LNCaP and DU145 transfected with the empty vector, EV (left) or with pBORIS-CMV6 (right) were subjected to digestion with McrBC (+) at 37°C for 16 hrs followed by PCR using *AR* promoter specific primers (see Figure 4B). The PCR products were then resolved on the agarose gel. The PCR bands were quantified using ImageJ software and the percentage of methylation at *AR* promoter after each transfection was calculated as explained in the Materials and Methods. A graph below represents the results of this analysis. **B.** Prostate cancer cells LNCaP and DU145 transfected with the empty vector, EV (left) or with pBORIS-CMV6 (right) were used for ChIP assays performed with the anti-BORIS antibodies. The ChIP DNA samples were analyzed by qPCR using primers for amplification of selected AR-CTSs (main sites, *AR* CTS-1 and *AR* CTS-2, and additional sites *AR* CTS- 4 and *AR* CTS-5). The DNA was analyzed for *AR* CTS-1 through to CTS-6 by qPCR using corresponding primers; all reactions were performed in triplicates and mean fold enrichment relative to the control ChIP experiment with no antibody (designated as 1.0) were calculated; error bars indicate the standard deviation (Student’s T-test: ** P<0.01; *** P<0.005).

### BORIS and CTCF interact differentially with the CTSs within the AR gene (AR CTSs) and the active state of the AR gene is associated with open chromatin configuration at the AR CTS-1 and AR CTS-2

To further analyze BORIS and CTCF binding within the predicted *AR* CTSs, ChIP assays were performed in two prostate cancer cell lines, LNCaP (ADPC, AR positive) and DU145 (AIPC, AR negative) using anti-BORIS and anti-CTCF antibodies. The qPCR amplification of the immunoprecipitated DNA reveals the enrichment by BORIS at the *AR* CTS-1 in both cell lines, although the levels were much higher in LNCaP cells than in DU145 cells. BORIS occupancy at other CTSs (*AR* CTS-2, -3, -4, -5 and -6) were considerably weaker compared to the *AR* CTS-1 in both cell lines (Figure 5A, left). In contrast to BORIS and in agreement with the ChIP-Seq data (Figure 4 and Supplemental Figure 6), binding of CTCF was lower at the *AR* CTS-1 and higher at the *AR* CTS-2 (Figure 5A, right). Increased binding by CTCF was also detected at the *AR* CTS-3 in both cell lines, and *AR* CTS-4 in DU145. Collectively, these data demonstrate that (i) CTCF and BORIS differentially bind to the human *AR* gene, (ii) their binding patterns are cell line specific and (iii) they may compete for binding to *AR* CTS-1 and -2, which are likely to be most important for the regulation of the *AR* gene due to their localization close to the *AR* gene transcription start site (TSS)/promoter.

To explore the association between the *AR* activity and the CTCF/BORIS binding further, the active (H3K4me3) and repressive (H3K27me3) chromatin marks at the *AR* CTSs in LNCaP and DU145 cells were examined. The *AR* CTS-1 and -2 were associated with active (open) chromatin marks in LNCaP, whereas the remaining four *AR* CTSs in LNCaP cells had a higher proportion of repressive (closed) chromatin. In DU145 cells, the *AR* CTS-1 and -4 were strongly associated with closed chromatin marks, whereas the fold enrichment by both marks at other *AR* CTSs was overall similar and relatively low (<10%).

From these experiments we conclude that BORIS preferentially binds to *AR* CTS-1 whereas CTCF binds to *AR* CTS-2 in both cell types. The active state of the *AR* gene in LNCaP (ADPC) cells is associated with higher occupancy by BORIS at the *AR* CTS-1 and open chromatin configuration at the *AR* CTS-1 and *AR* CTS-2, whereas the inactive state of the *AR* gene in DU145 (AIPC) is associated with higher occupancy by CTCF at the AR CTS-2 and closed chromatin at all *AR* CTSs.

### Introduction of ectopic BORIS leads to DNA hypomethylation within the AR gene promoter

BORIS in cancer cells may play a role in the epigenetic regulation by maintaining the hypomethylated status of promoter regions of some oncogenes [15, 22]. To investigate a possible link between BORIS and the methylation status of the *AR* promoter, DNA from LNCaP (ADPC) and DU145 (AIPC) cell over-expressing BORIS was digested with a methylation dependent restriction enzyme, McrBC. This endonuclease specifically cleaves DNA containing methyl cytosine preceded by a purine nucleotide (A or G) and as a result, methylated DNA, treated with McrBC can not be amplified. In agreement with published observations [67, 70], in control cells transfected with the empty vector, high levels of methylation (92%) was observed in the *AR* promoter region in DU145 cells, in comparison with 17% in LNCaP cells (Figure 6A). In both cells over-expressing BORIS the *AR* promoter and almost fully de-methylated in LNCaP cells and 32% methylated in DU145 cells. More bound BORIS protein was found at the *AR* CTS-1 and *AR* CTS-2 in cells transfected with the pCMV-BORIS in both cell lines (Figure 6B). This phenomenon is more pronounced at the *AR* CTS-2 characterized by higher occupancy by CTCF in untreated cells indicating that BORIS may outcompete CTCF at this site.

### Over-expression of BORIS reduces CTCF binding and repressive chromatin marks, and increases active chromatin marks within the AR gene

Our observations so far indicated that low occupancy by BORIS at the *AR* CTS-1 in AIPC (DU145) cells may be linked to DNA hypermethylation and enrichment with repressive chromatin, hereby leading to *AR* gene silencing in these cells. As introduction of BORIS into these cells resulted in re-activation of *AR*, we hypothesized that at increased concentrations BORIS can outcompete CTCF and change the chromatin state of the *AR* gene, in particular at the promoter region. To investigate this, prostate cancer cell lines LNCaP and DU145 were transfected with a DNA vector containing BORIS cDNA followed by investigation of CTCF occupancies and chromatin states at the *AR* CTSs by ChIP (Figure 7A and B). Presence of the ectopic BORIS protein in transfected cells was confirmed by Western blot analyses (Figure 7C). The binding of CTCF in both cell lines was generally higher in control cells transfected with the empty vector and lowest in cells transfected with pCMV6-BORIS. These findings indicate that over expression of BORIS in prostate cancer cells can displace CTCF from its target sites, which may have implications on the AR expression and prostate cell proliferation.

**Figure 7.**
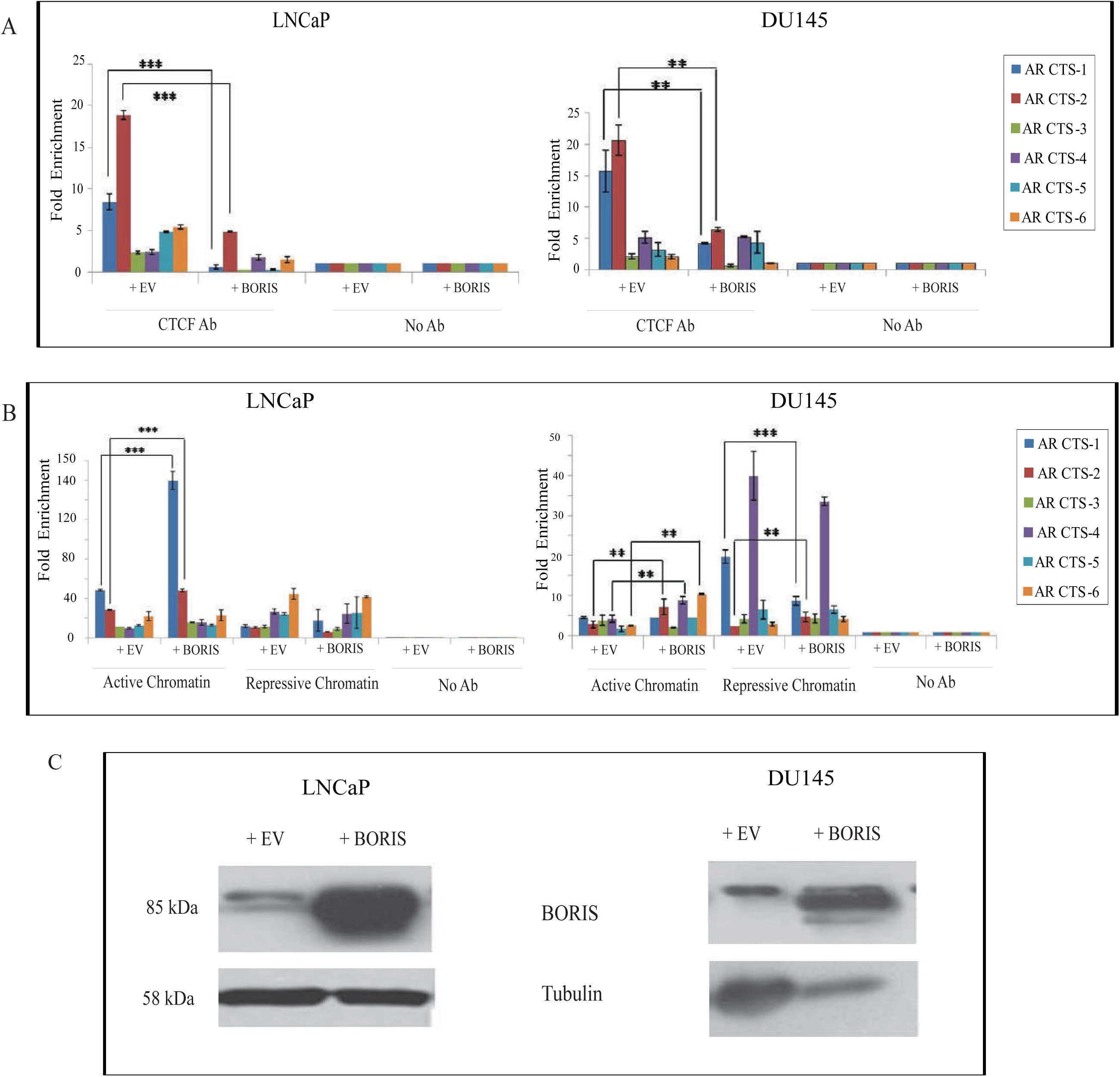
Analysis of CTCF binding and distribution of chromatin marks and the *AR* promoter methylation at the *AR* CTSs. LNCaP and DU145 cells transfected with empty vector (+EV) and pCMV6-BORIS (+BORIS), ChIP essays were performed, ChIP reaction was carried out using anti-CTCF, anti-H3K4me3 (active chromatin) and anti-H3K27me3 (repressive chromatin) antibodies. The DNA was analyzed for *AR* CTS-1 through to CTS-6 by qPCR using corresponding primers; all reactions were performed in triplicates and mean fold enrichment relative to the control ChIP experiment with no antibody (designated as 1.0) were calculated; error bars indicate the standard deviation (Student’s T-test: ** P<0.01; *** P<0.005). **A.** Analysis of CTCF binding in transfected LNCaP/DU145 cells. **B.** Analysis of chromatin marks in transfected LNCaP/DU145 cells. **C.** Western blot analyses of LNCaP and DU145 cells transfected with empty vector (+EV) and pCMV6-BORIS (+BORIS) used for ChIP; α-tubulin levels were used as the loading control.

The transfected LNCaP and DU145 cells were also used to investigate whether introduction of CTCF and BORIS into cells could affect the chromatin states at the *AR* CTSs. To achieve this, ChIP assays were performed using the antibodies for anti-H3K4me3 (active chromatin mark) and anti-H3K27me3 (repressive chromatin mark). The results of qPCR analyses of ChIP DNA are presented in Figure 7B. In comparison with the control cells transfected with empty vector, in LNCaP (ADPC), over-expressing BORIS, considerable enrichment with open chromatin marks was observed at *AR* CTS-1 and *AR* CTS-2, whereas in cells over-expressing CTCF all *AR* CTSs were enriched with repressive chromatin marks. Interestingly, in DU145 (AIPC) increased levels of BORIS resulted in modest but notable elevation of active chromatin marks at all *AR* CTSs except CTS-3. On the other hand, the presence of repressive chromatin marks decreased at the *AR* CTS-1 and *AR* CTS-4 in these cells. Over-expression of CTCF in DU145 cells did not change the level of active chromatin marks, however it led to considerable increase of repressive chromatin marks at all *AR* CTSs.

In summary, the experiments described in this section demonstrate that elevation of BORIS levels results in the reduction of CTCF binding, increase of active and decrease of repressive chromatin marks at the *AR* CTS-1 and CTS-2 in both LNCaP (ADPC) and DU145 (AIPC) cells. These data suggest that BORIS and CTCF may be able to remodel the chromatin within the *AR* gene and play a role in the regulation of *AR* at the transcriptional level.

## Discussion

In this study we aimed to investigate a role of BORIS in regulating the *Androgen Receptor (AR)*, one of the major genes vital for normal prostate function and for the initiation of prostate cancer development. Tumours, initially dependent on androgens (ADPCs) slowly progress to more aggressive androgen independent prostate cancers (AIPCs). Evidence exist that a modest increase in *AR* mRNA is the only change consistently associated with this progression and the development of resistance to anti-androgen therapy [71, 72]. BORIS is likely to play a key role in these processes due to its ability to positively regulate the *AR* gene expression (this report), in addition to correlation between higher levels of BORIS with increased AR and more aggressive disease described in our previous study[18].

A wide range of genetic and epigenetic mechanisms have been implicated in the development of the AIPCs [48-50], negative for *AR* mRNA and protein in some tumours but increased in others [73].Epigenetic changes are of particular interest as they can be reversed and cells then may restore their sensitivity to anti-androgens if the normal *AR* gene is silenced. Indeed, treatment with demethylating chemicals, such as 5-aza-20-deoxicytidine, has been successful in partial re-activation of *AR* in AIPC cells[67]. A known association of BORIS with the re-setting of genome methylation during early embryonic male germ cell development[1, 2] and BORIS ability to reactivate various genes[22-24] [25-27] prompted us to investigate BORIS role in the regulation of the *AR* gene in ADPC and AIPC cells.

Following the introduction of the ectopic BORIS, we observed AR over-expression in ADPC (LNCaP and VCap) cells and re-activation of AR in AIPC cells (DU145 and PC3), both at the mRNA and protein levels. Depletion of ADPC cells of BORIS led to reduction of AR; no change in AR was detected in AIPC cells negative for AR in basal conditions (Figures 1 and 3). These findings indicate that regulation of AR expression by BORIS takes place at the level of transcription, which is not surprising given that BORIS has features characteristic for a transcription factor and has been demonstrated to regulate transcription of other genes[1, 17, 21]. In our experiments, AR was found to be negatively regulated by CTCF, although this effect was demonstrated only in CTCF knock down experiments (Figure 1). Taken together, these findings indicated that there may be functional interplay between CTCF and BORIS in the regulation of AR, which is further discussed below. In general, we noted the lack of correlation between the *AR* mRNA and the AR protein levels for all three mRNA/protein pairs (AR, CTCF and BORIS) (Figure 1). This phenomenon may be explained by differential stability of mRNA and proteins, which may also depend on cell context, with broader variation for proteins than for mRNA, whereby the half-life of different proteins spans from minutes to days compared to 2-7 hours for mRNA [74].

The AR produced in AIPC cells over-expressing BORIS was biologically active by several criteria: sensitivity to DHT, translocation into the nucleus after treatment with DHT, and activation of the AR-dependent genes, *PSA* and *TMPRSS2*. The AR produced in DU145 cells containing ectopic BORIS-EGFP was translocated to the nucleus of these cells after DHT treatment, but not in the absence of DHT, whereas control cells with ectopic EGFP did not express AR, with or without DHT. Similar effects were observed in another AIPC cell line, PC3, thus further supporting this phenomenon (Figure 2). As expected, AR was detected in the nucleus of ADPC (LNCaP) cells in the presence of DHT, however significantly higher levels of AR were observed in these cells in the presence of DHT and ectopic BORIS-EGFP. Up-regulation of AR by BORIS, together with AR autoregulation[75], may saturate the amount of AR in the nucleus and inhibit further translocation of AR. Similar observations in LNCaP cells were reported demonstrating that very high levels of AR can be lethal for the cells and negative feedback AR regulation may be important to control cell growth and avoid transition to androgen independence [76, 77]. The latter may also explain our results showing that levels of *PSA* and *TMPRSS2* mRNA in ADPC (LNCaP and VCaP) cells containing ectopic BORIS and DHT were similar to the respective control cells with the empty vector. Induction of *PSA* and *TMPRSS2* mRNAs in AIPC cells comparable with these levels was only achieved in cells containing DHT and ectopic BORIS, which suggests the activation of appropriate upstream signalling pathways involving AR. We therefore expect that by restoring the sensitivity of the AIPC cells to androgens other cellular functions will be properly regulated by the *de novo* produced AR protein. Introduction of the ectopic BORIS, with and without DHT treatment, had no effect on *PSA* and *TMPRSS2* in BPH-1 cells (Figure 3) and *AR* expression (data not shown), suggesting a different mechanism of the *AR* silencing, which may be due to genetic changes (e.g. absence of *AR* or a large deletion in the gene) or reflect a possible BPH-1 origin from AR- negative prostate stem cells [78].

Since BORIS shares the same DNA binding domain with CTCF, we reasoned that CTCF binding sites (*AR* CTSs) identified within the AR gene using bioinformatics analysis of the ChIP-Seq (Figure 4) may also bind BORIS. Five of the identified six *AR* CTSs are positioned in introns, whereas *AR* CTS-1 is localized in exon 1. Such intragenic positioning of the CTSs is not surprising given that a large proportion of CTSs are reported in gene regions other than promoter and 5’ UTRs [69, 79]. Furthermore, the concept of the spread of regulatory elements throughout the genome regions, intra- and intergenic, has been generally accepted [80, 81].

ChIP analyses of the *AR* CTSs revealed differential binding of CTCF and BORIS, with the *AR* CTS-1 and *AR* CTS-2 as main sites. Although no negative control of a non-CTS region is included in the ChIP experiments, some of the putative CTS sites (*AR* CTS-5 and *AR* CTS-6) act as control sites where no CTCF or BORIS binding is observed (Figure 5). Interestingly, in both ADPC and AIPC cells, CTCF was predominantly bound to *AR* CTS-2 and less to *AR* CTS-1, whereas *AR* CTS-1 was the main site for BORIS binding (Figure 5A). Both sites are located downstream of the transcription start site, with the *AR* CTS-1 spanning the end of 5’UTR and continuing into exon 1,whereas *AR* CTS-2 is positioned in intron 1. As reported previously, such position of CTSs is characteristic for genes negatively regulated by CTCF [5, 82, 83].

The analysis of BORIS, CTCF and active/repressive chromatin at the *AR* CTSs revealed the link between BORIS and active state of the *AR*. The *AR* CTSs were more enriched with BORIS and active chromatin marks in ADPC (LNCaP), compared to AIPC (DU145) cells, On the other hand, the enrichment with CTCF and repressive chromatin marks at the *AR* CTSs was evident in DU145 cells(Figure 5B). It is possible that the presence of CTCF may facilitate the recruitment of enzymes converting DNA into highly condensed chromatin thus making it inaccessible to other transcription factors or co-factors necessary for *AR* expression. Indeed, CTCF was previously reported to recruit histone deacetylases [84] and Polycomb repressive complex 2 (through its interaction with Suz 12) [85], leading to transcriptional repression.

The presence of ectopic BORIS correlated with the reduction of methylation in the *AR* promoter in both ADPC and AIPC cells (Figure 6). Such reversible epigenetic changes (hypo-or hyper-methylation) affecting gene regulation are common in cancer cells [86]. Studies using haematological and solid tumours have reported a small subset of dedifferentiated cells that are capable of developing stem cell phenotype; they were more likely to be associated with DNA hypomethylation [87, 88]. Interestingly, expression of BORIS has been associated with proliferation and erasure of methylation marks during male germ-line development [2] and with hypomethylation in cell lines and tumours [22, 89].

Our findings suggest the importance of BORIS in the epigenetic regulation of *AR* function in prostate cells, summarized in the model presented in Figure 8. In ADPC, BORIS preferentially binds the *AR* CTS-1 whereas CTCF is mainly bound to the *AR* CTS-2 and this configuration leads to the establishment of open chromatin, hypomethylation of DNA and transcription of the *AR* mRNA (Panel A). In AIPC, CTCF is predominantly bound to *AR* CTS-2 and very low or no BORIS binding is observed at the *AR* CTS-1; such distribution of CTCF and BORIS is associated with the repressive state of the chromatin, DNA methylation and silencing of *AR* (Panel B). Following the introduction of the ectopic BORIS, its occupancies at the *AR* CTS-1 and *AR* CTS-2 increase in both, AIPC and ADPC, cells. This is associated with the reduction of CTCF binding, especially in AIPC cells, which is likely to be due to out-competition of CTCF by BORIS, increase with the marks characteristic for open chromatin, DNA hypomethylation and increase in the production in ADPC cells (Panel A) or re- activation of the *AR* mRNA in AIPC cells (Panel B). Of note, although the ectopic BORIS in ADPC cells increases the levels of *AR*, it is not dramatic and may be explained by a possible feedback mechanism restricting uncontrollable expression of *AR* which may be detrimental for ADPC cells (Panel A).

**Figure 8.**
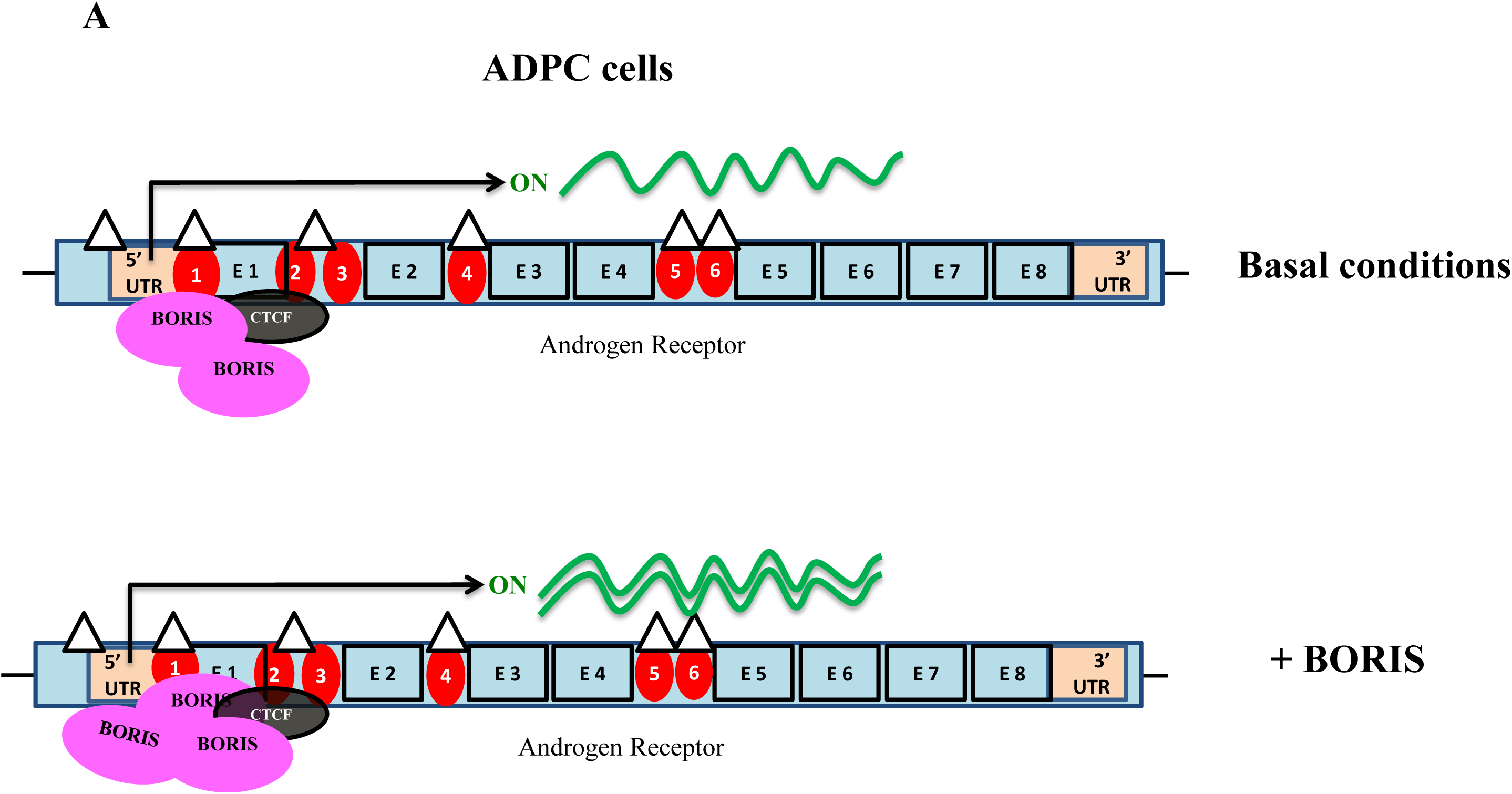
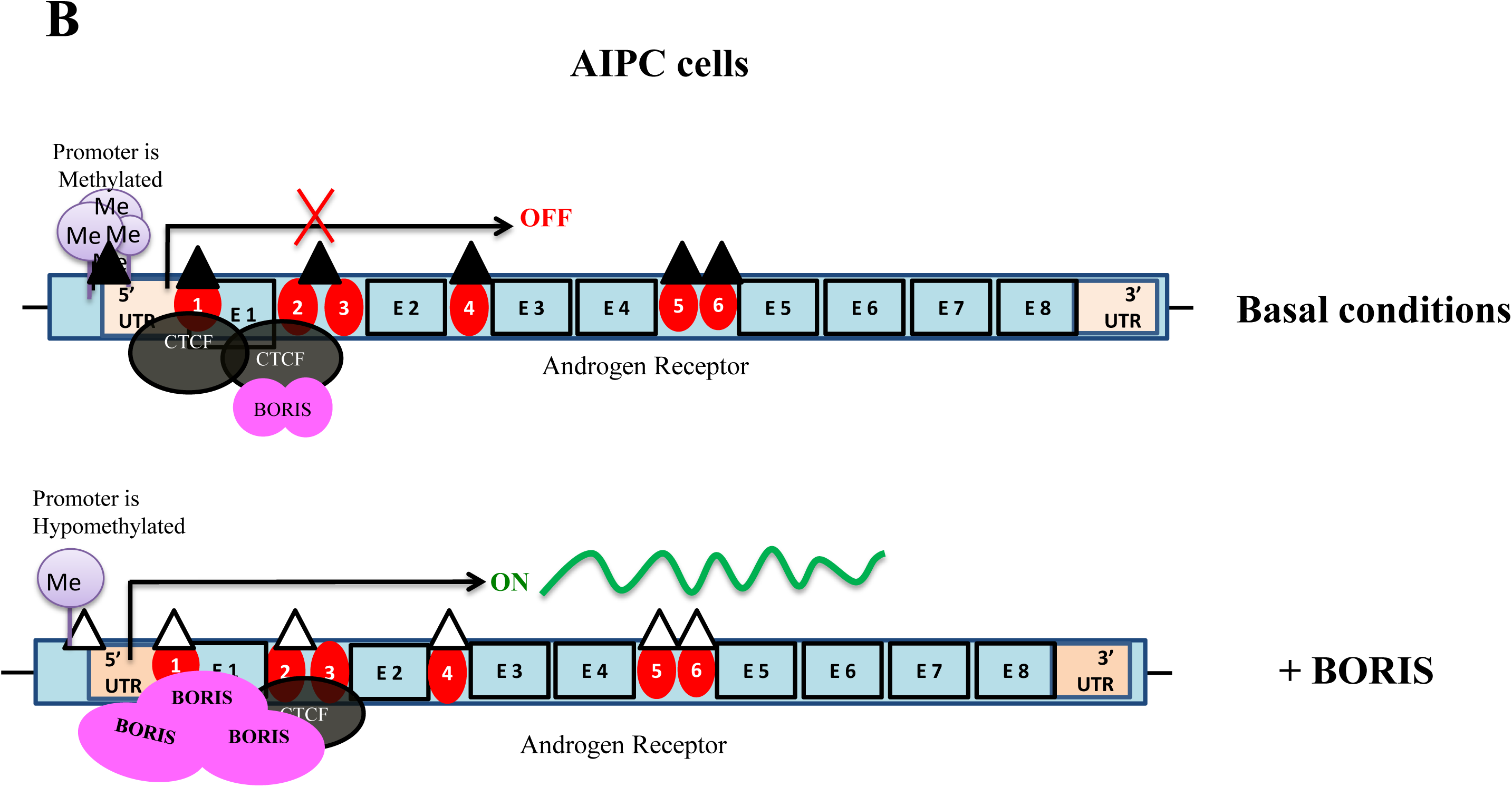
Model for epigenetic regulation of *AR* by BORIS and CTCF in ADPC and AIPC cells. **A.** *AR* regulation by BORIS in ADPC in basal conditions and in cells-over expressing BORIS. In ADPC cells with active *AR* expression BORIS is preferentially bound to *AR* CTS-1, and the upstream *AR* CTSs sites are enriched in open chromatin marks. Ectopic BORIS expression in ADPC leads to over expression of AR however production is saturated after a certain level. Panel **B:** *AR* regulation by BORIS in AIPC in basal conditions and in cells-over expressing BORIS. In AIPC, the silenced state of AR is associated with reduced occupancy of the *AR* CTS-1 by BORIS and increased binding of *AR* CTS-2 by CTCF, promoter hypermethylation and closed chromatin marks. At higher levels BORIS outcompetes CTCF at *AR* CTS-1 and *AR* CTS-1, leading to erasure of methylation marks, establishment of active chromatin and activation of transcription from the *AR* gene. Pink circles: BORIS; Black circles: CTCF; Empty triangles: open chromatin marks; Filled triangles: closed chromatin marks.

The direct competition between CTCF and BORIS for binding to the *AR* CTSs may explain the changes in the *AR* activity (Figure 8), however the *AR* regulation by CTCF/BORIS may be more complex and involve other transcription factors regulated by CTCF/BORIS including negative effects of CTCF on *BORIS* expression [21, 90]. Since AR is associated with the development of more aggressive prostate cancer [45], BORIS function as the transcriptional activator of *AR* would generally support tumour progression, whereas CTCF which inhibits growth of prostate cancer cells can counteract such influences. These antagonistic features of BORIS and CTCF have been reported previously, however this mode of function may be limited to a certain subpopulation of the DNA targets [13] and the *AR* gene is likely to belong to this category. It should be noted that the ability of BORIS to activate *AR* is likely to be restricted to certain stages in the prostate tumour progression. When the androgen-independent state is established in the process of tumour evolution, the levels of endogenous BORIS may not be sufficient to activate the epigenetically silenced AR gene. However, re-activation of *AR* may be possible if BORIS levels in the cells increase. In the context of this model, stimulation of BORIS expression can be used to re-sensitize AIPC cells so that they can be treated with anti-androgens. It needs to be mentioned that the proposed model is mostly relevant to a subgroup of the AIPC in which the *AR* gene is epigenetically silenced. It is therefore important to determine the molecular basis of the androgen independency to design appropriate treatment strategies. Finally, whereas this study was focused on the *AR* as a specific regulatory target of BORIS in prostate cells, it should be acknowledged that as a global transcriptional factor BORIS has wider effects on cellular processes, in particular supporting signalling pathways associated with cell survival and proliferation. The genome-wide expression analysis of prostate cancer cells with manipulated levels of BORIS and CTCF will be published in a separate article (Y.Hari-Gupta, manuscript in preparation).

In conclusion, this study provides novel insights into the role of BORIS in the regulation of the *AR* gene, which may lead to better understanding of the mechanisms of prostate tumourigenesis and, potentially, development of clinical applications.

## Acknowledgements

We are grateful to S. Hayward for providing the BPH-1 prostate cell lines and V. Lobanenkov, D. Loukinov and D. Delgado for providing us with plasmids pCMV-BORIS and pEGFP-BORIS. We thank E. Pugacheva for assistance with the bioinformatics analysis and A. Angel for excellent technical assistance. We also thank members of Elena Klenova’s, Vladimir Teif’s and Greg Brooke’s laboratories for experimental advice and many helpful discussions.

### Funding

This work was supported by the Colchester Catalyst Charity (EK and YH-G, PhD studentship), Cancer Research UK (C8787/A12723, EK and DF), Medical Research Council (G0401088, EK and DF); Breast Cancer Research Trust (EK and G-XK), Breast Cancer Campaign (2010NovSP22, EK and G-XK), University of Essex (EK and G-XK).

